# A network approach to genetic circuit designs

**DOI:** 10.1101/2021.09.14.460206

**Authors:** Matthew Crowther, Anil Wipat, Ángel Goñi-Moreno

## Abstract

As genetic circuits become more sophisticated, the size and complexity of data about their designs increases. This data captured goes beyond monolithic genetic sequences and towards circuit modularity and functional details, which are beneficial for analyzing circuit performance and establishing design automation techniques. However, the accessibility, visualisation and usability of design data (and metadata) have received relatively little attention to date. Here, we present a method to turn circuit designs into networks and showcase its potential to enhance the utility of design data. Since networks are dynamic structures, initial graphs can be interactively shaped into sub-networks of relevant information based on requirements such as abstraction, hierarchy and protein interactions. Additionally, several visual changes can be applied, such as colouring or clustering nodes based on types (e.g., genes or promoters), resulting in easier comprehension from a user perspective. This approach allows circuit designs to be coupled to other networks, such as metabolic pathways or implementation protocols captured in graph-like formats. Therefore, we advocate using networks to structure, access and improve synthetic biology information.

## Introduction

The design and implementation of genetic circuits^1,2^ that allow cells to perform predefined functions lies at the core of synthetic biology.^3,4^ For example, the engineering of Boolean logic circuits^5^ that use cascades of transcriptional regulators is a field that regularly scales up both complexity and functionality. Other types of circuits are routinely engineered, such as switches,^6^ counters^7^ and memories;^8^ using not only transcriptional, but also post-transcriptional, processes.^9^ Different host organisms such as bacteria,^10^ yeasts^11^ and mammalian^12^ cells are used to test circuits in several applications, ranging from pollution control^13^ to medical diagnosis.^14^ Furthermore, the functionalities of genetic circuits will only improve as scientists control the information processing abilities of living substrates: signal noise,^15,16^ metabolic dynamics,^17,18^ context-circuit interplay,^19,20^ stability^21^ and more.^22^

Due to the cyclical nature of synthetic biology projects combined with complexity and size, successful implementation and testing can be challenging without a well-conceived design and solid understanding. Mathematical and computational tools,^23^ automation methods,^24,25^ knowledge-based systems^26,27^ and repositories^28^ assist circuit design to minimise the iterations within the design-build-test-learn life-cycle. These processes generate a consortium of information beyond unitary DNA sequences, such as modularity, hierarchy, implementation instructions, dynamical predictions and validation strategies. However, this information is often overlooked, which threatens to undermine the success of such endeavours.

Data formats have emerged that effectively capture and represent increasingly complex designs. A leading example is the development of the Synthetic Biology Open Language^29^ (SBOL) that can be used to represent both structural (e.g. DNA sequences) and functional (e.g. regulation interactions) information. Also, the GenBank^30^ format, overwhelmingly used to formalize and share genetic sequences, allows simple annotations to be defined. The overarching challenge is to access, use, visualize and analyse this information so that genetic designs become dynamic data structures easy to manipulate. This challenge underpins this work, where we propose networks to help solve these problems.

What has been termed *network biology*^31^ deals with the quantifiable representation of complex cellular systems in graphs and the study of them in order to characterise functional behaviour. Indeed, the analysis of interaction maps (a type of network) can reveal previously unknown mechanistic details. Graph theory methods can assist the interrogation of network structures in several ways^32^ for circuit designs, where dynamic querying produce sub-networks of particular interest hidden within design formats. In order to build networks out of circuit designs, data is represented in the form of nodes (individual points of data) and edges (relationships between the data).^33^ For example, a repression relationship edge would link two nodes representing a regulator protein (e.g. aTc) and its cognate promoter (pTet). The relevance of this approach is enhanced by the emergence of high-throughput techniques and workflows^34,35^ that generate (and use) large data sets.

Visualising complex information is a challenge within synthetic biology mainly because a one-size-fits-all approach is often not feasible, i.e. a single representation of a multidimensional dataset cannot satisfy the requirements of all involved members. For example, take the glyph approach,^36^ where each genetic part is displayed on a linear sequence. While this allows researchers to generate diagrams to visualize and communicate abstract designs, an experimentalist cannot access information regarding nuanced sequence data, or an informatician cannot explore the provenance of a specific genetic part. In contrast, the presented network visualisations can be dynamically adjusted according to specific requirements, such as highlighting proteins, interactions or hierarchy. Here, we demonstrate how network techniques can be applied to the analysis of genetic circuit design data. While the focus is primarily on displaying designs into comprehensible visualisations, many graph-based approaches are combined to produce the final visualized output.

## Results and discussion

### Establishing networks from design files

Figure 1 shows the process of visualising and structuring the data encoded within a genetic circuit design. We used the design of the NOR logic gate built by Tamsir and colleagues.^37^ This gate outputs 1 (i.e., target gene expressed) if both inputs are 0, and outputs 0 (i.e., target gene not expressed) in any other case. The NOR circuit is a frequently built device,^5,38^ since any logic function can be achieved by assembling NOR gates only.

**Figure 1:**
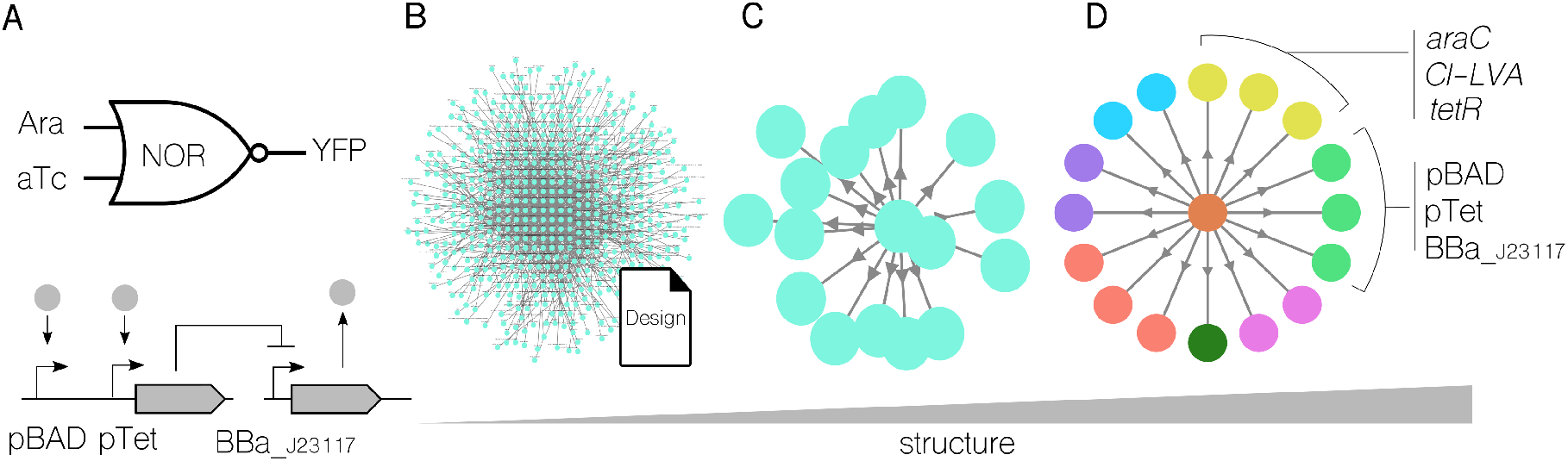
Visualising design data of a NOR logic gate. **A**. NOR logic function and genetic diagram, with inputs (arabinose; aTc) and output (YFP). **B**. Displaying all design information (including metadata) in network format. The network is unreadable but computationally tractable. **C**. A network is generated from the same design where only the physical elements (i.e. DNA and molecular entities described) are shown. **D**. Depending on their role, the network is adjusted to display colours for the nodes for visualisation purposes. Roles (e.g. promoter, proteins) are automatically clustered by the same colour. For clarity purposes, labels were not included.

The functional diagram for the NOR gate (Figure 1A top) is often represented with the specific names of the input and output compounds. In this case, the inputs are the inducers arabinose (Ara) and anhydrotetracycline (aTc), and the output reporter yellow fluorescent protein (YFP). The genetic visualisation (Figure 1A bottom) is usually labelled with names for specific DNA parts, three promoters in this example. While these representations serve a purpose, they offer only single insights into larger data structures. Figure 1B shows the results of building a network with all the available data and metadata. Although this network is visually meaningless because the significance of nodes and connections is lost, the graph is established, and computational manipulation is easy to perform based on specific requirements.^39^ The network of Figure 1C is a *view* into the data that focuses on physical elements only (e.g., DNA and molecular entities) and omits metadata details. By presenting a single perspective, overall cognition is increased, but it is visually incoherent since the position of nodes–its *layout*—is random. The layout, which determines the arrangement of nodes combined with other features such as colour, size and shape,^40^ provides a visual representation of information that ensures clarity and understanding. When a simple radial layout (Figure 1D) (e.g., nodes do not overlap) is combined with node clustering and colouring depending on types and roles, it results in a final visual output that is considerably clearer than fig:Figure 1B.

### Dynamic abstraction levels

The design of a biological system implies dealing with complexity. Therefore, it is crucial to abstract away superfluous details to describe and communicate the design with clarity.^41^ Nevertheless, what is an appropriate level of abstraction for a circuit design? The answer depends on two primary factors: what information needs to be communicated, e.g., structural or functional, and the requirements of the person consuming the information, e.g. Bioinformatician or *wet lab* scientist. Precisely, a vital advantage of a network structure is its inherent ability to arrange itself—dynamically—into several levels of abstraction.

Interaction networks provide a high-level metric with the potential to scale, and this view into the data has been chosen to display dynamic abstraction. Figure 2 shows an interaction network of a NOR gate design at three different levels of abstraction. While the design is, of course, the same—that is, the file is not modified—the resulting network can differ depending on user needs. The network in Figure 2A displays all molecular, genetic, element types and relational information. Half of that information is encoded into the colour scheme of the graph; for example, node *yfp* is a coding sequence (yellow) that leads to the production of (yellow edge) YFP, which is a protein (red node). The arrangement of the nodes is far from random; it helps transmit information concerning the flow of information. In this case, information flows from inputs (upper nodes) to outputs (lower node). This network can be adjusted to remove all non-genetic elements (Figure 2B), limiting the information to only the DNA sequences described in the design (e.g., promoters and genes). Even in this case, relational information remains; for instance, the coding sequence *araC* represses the promoter node BBa_.J23117. Here, the abstraction level hides specific mechanistic information about the regulatory protein that performs repression and implicit activation. In order to increase abstraction by reducing superfluous connections, we used graph theory methods and transitivity-based algorithms (see Methods) that estimate the costs of different paths across the network to cluster related nodes,^42^ leading to a lower detailed but more straightforward network. Finally, the highest level of abstraction is to show input and output elements only ((Figure 2C), which allows for quick communication of circuit performance while abstracting away all implementation details and internal workings. This over-simplification may be excessive for a relatively simple design but could benefit more extensive and complex structures.

**Figure 2:**
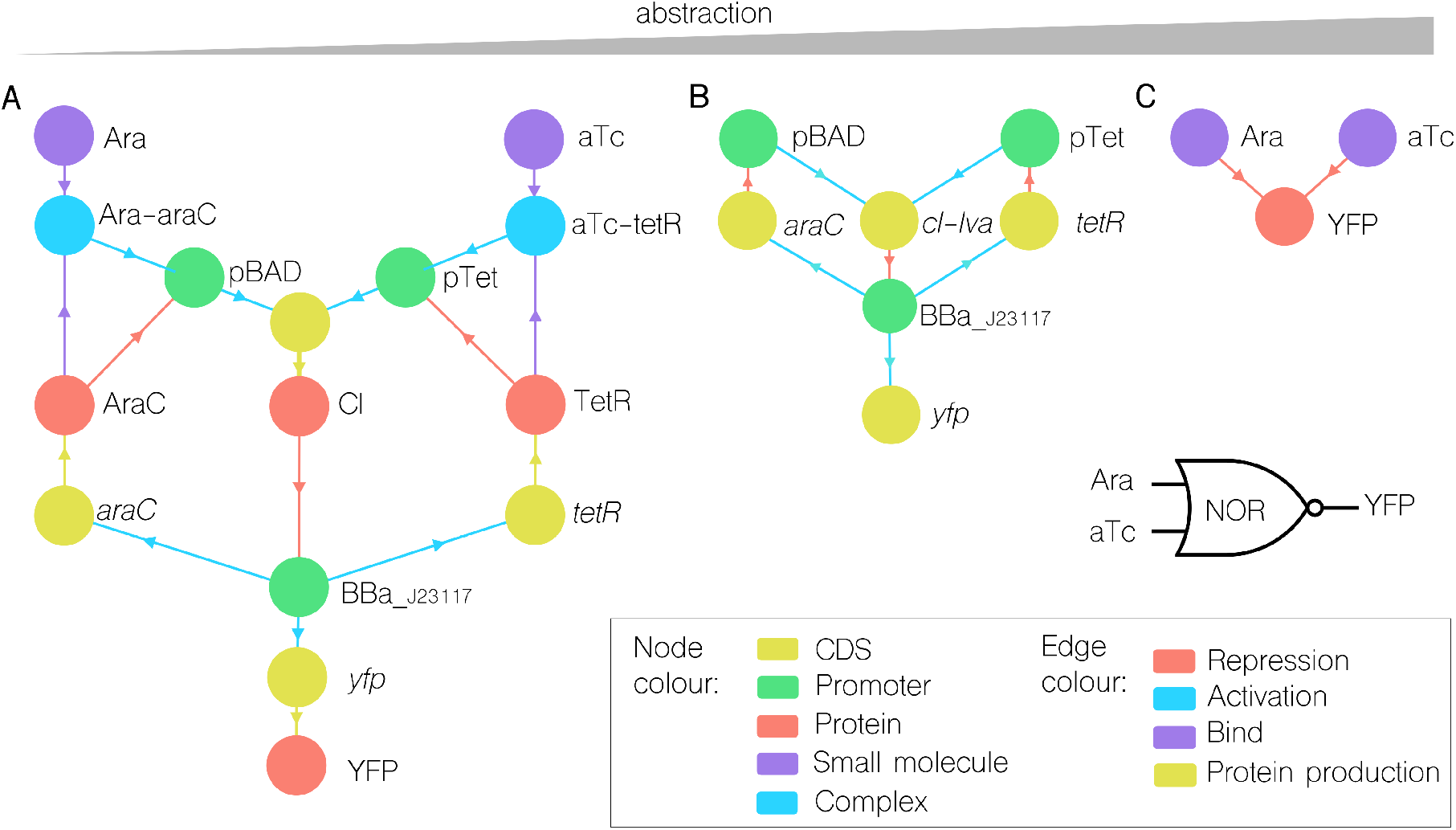
Adjusting network abstraction levels. **A**. The NOR gate design is turned into a network with all molecular and genetic elements (nodes); and interactions between entities (edges). **B**. Path-based analysis over the initial graph clusters nodes within the same hierarchical path, reducing complexity while retaining information flows. For instance, Ara, Ara-araC and pBAD are simplified into a single pBAD node. **C**. Maximum abstraction into input-output data. The colour scheme is constant regardless of abstraction levels.

### Hierarchical trees

Structuring genetic parts into a hierarchical tree of modules^44^ is crucial for scaling up circuit complexity and one of the design principles to turn biology into an engineering discipline. A *tree* is a fundamental network topology that can display a hierarchical representation of arbitrary depth. Different abstraction layers can be visualized simultaneously–the top node with the highest abstraction level and the bottom nodes where most details are described. As the size of genetic circuits increases, hierarchical networks facilitate the structural organization of information.

Figure 3 shows the hierarchical network corresponding to the *digitalizer* genetic circuit.^43^ This device is far more complex than the previous NOR logic gate, both in terms of interactions and dynamic performance, therefore ideal for showcasing the use of hierarchical representation. The goal of the *digitalizer* circuit is to minimize the leakage expression of a specific gene of interest while maximizing the full production. That is to say, to enlarge its dynamic range. The circuit (Figure 3A) is based on two negative interactions, between the regulatory protein LacI and a small RNA, and offers the ability to plug and play any gene of interest the user wants to *digitalize—*the reporter *gfp* gene is used for characterization. Its hierarchical tree (Figure 3B), which is automatically built from the design file, displays the conceptual modules into which single parts are structured. This information is often particular for each circuit-even similar or identical circuits-since it follows the authors’ conceptual framework. In this case, the top module represents the whole device and is broken down into four modules, which are, in turn, leading either to final parts (e.g., promoters *Pm* and *P-A1/04S*) or to smaller sub-modules (e.g., GFP cassette).

**Figure 3:**
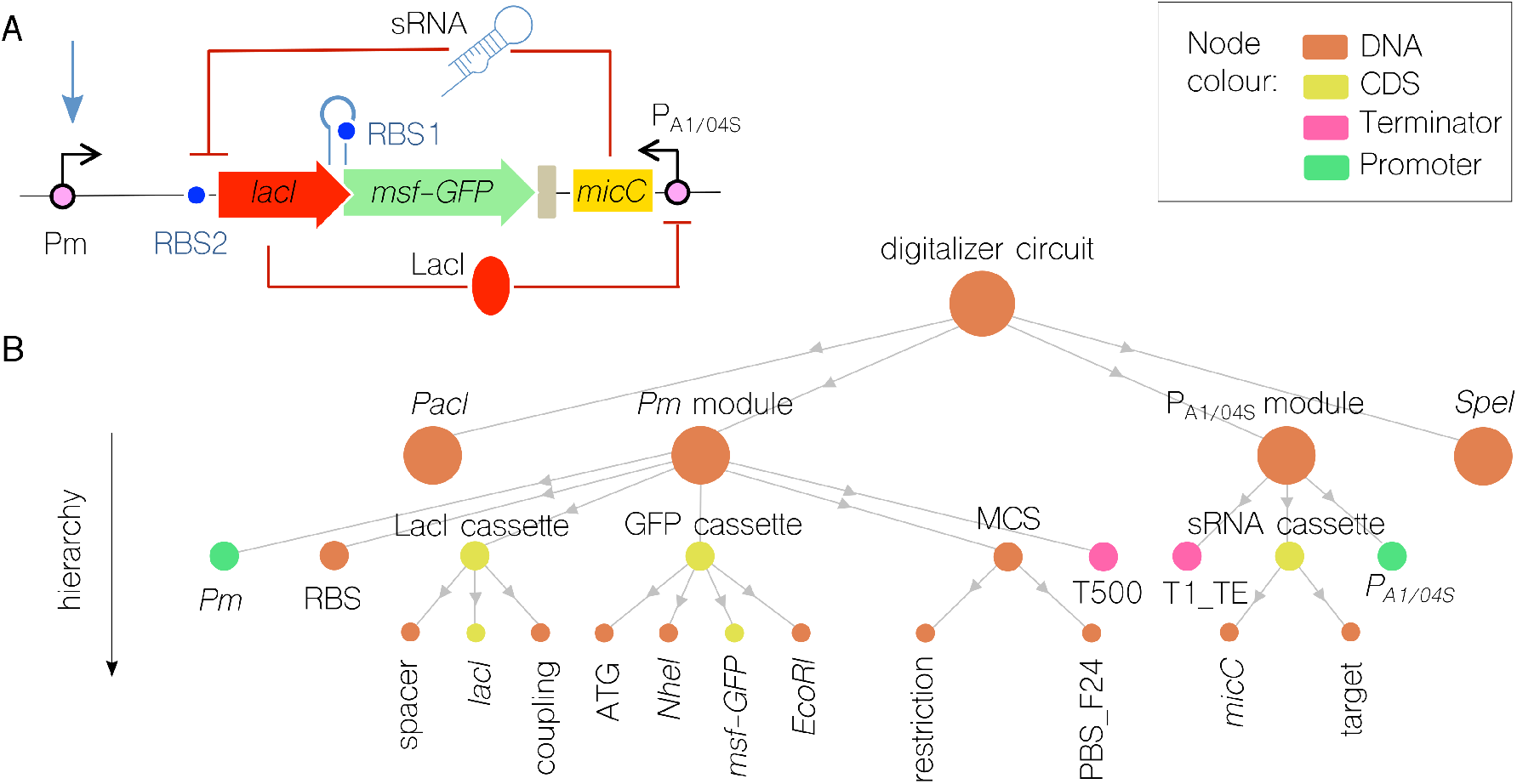
A hierarchical network of increasing abstraction, from modules to parts. **A**. The *digitalizer*^43^ synthetic circuit showcases the hierarchical information encoded within the design file. **B**. The resulting network is where bottom nodes are single DNA parts, and nodes in higher levels represent modules, the top node being the entire circuit. Circuit building details are highlighted within the network, e.g. restriction sites or sequence to couple *lacI* to *msf-GFP*. Functional details are also displayed as the strategy to change the *msf-GFP* gene using *NheI* and *EcoRI* restriction sites. Edges represent hierarchical direction.

Specific structural details that refer to implementation strategies are essential in those genetic circuits whose goal is to let users modify parts of them. The *digitalizer* circuit is an example where the user is meant to switch the reporter gene by his/her gene of choice. By browsing through the network in Figure 3B, the user can find a module where the reporter is included (named GFP cassette) and the hard-coded procedure for cutting out the gene (i.e., restriction sites *NheI* and *EcoRI*) without looking at the genetic sequence of the design.

### Protein interaction maps for representing the function

The interaction between its proteins better describes the functional performance of a genetic circuit than by the structural details of its DNA sequence. Therefore it is essential to complement sequence-based designs and visualisations with regulatory information. Interaction networks can provide a higher-level understanding by visualising regulatory proteins and abstracting relationships. For example, the mechanisms that allow a regulator to bind its cognate promoter and repress a downstream gene’s expression into another regulator can be abstracted away into a simple network with two nodes (one per regulatory protein) linked by an arrow. While this network lacks structural information at the sequence level (e.g., implementation details), it maximises the functional aspect of the circuit.

Figure 4 shows the protein interaction map of a circuit consisting of 5 sequentially connected logic gates (4 NOR and 1 OR) that builds a 3-input 1-output circuit. The sequence-oriented diagram (Figure 4A bottom) offers limited information. For example, it shows the number of promoters and genes required to synthesise (or assemble) the circuit, but it says little about its function. The whole graph generated from this circuit design is shown in Figure 4; while not displaying helpful information, it gives an idea of how much data the design has. From that data, a new sub-network with only regulators as nodes and their relationships as edges can be easily generated to display the protein interaction map of the circuit (Figure 4C). In the resulting network, the three input regulators (LacI, TetR and AraC) and the output protein (YFP) are clear, and information flows are explicitly displayed. Network topology also makes visualisation easier since nodes are arranged following functional criteria rather than sequentially. Moreover, the Boolean logic of the network becomes apparent; for example, the network contains a final OR logic gate based on the regulators PhiF or BetI repressing the presence of YFP.

**Figure 4:**
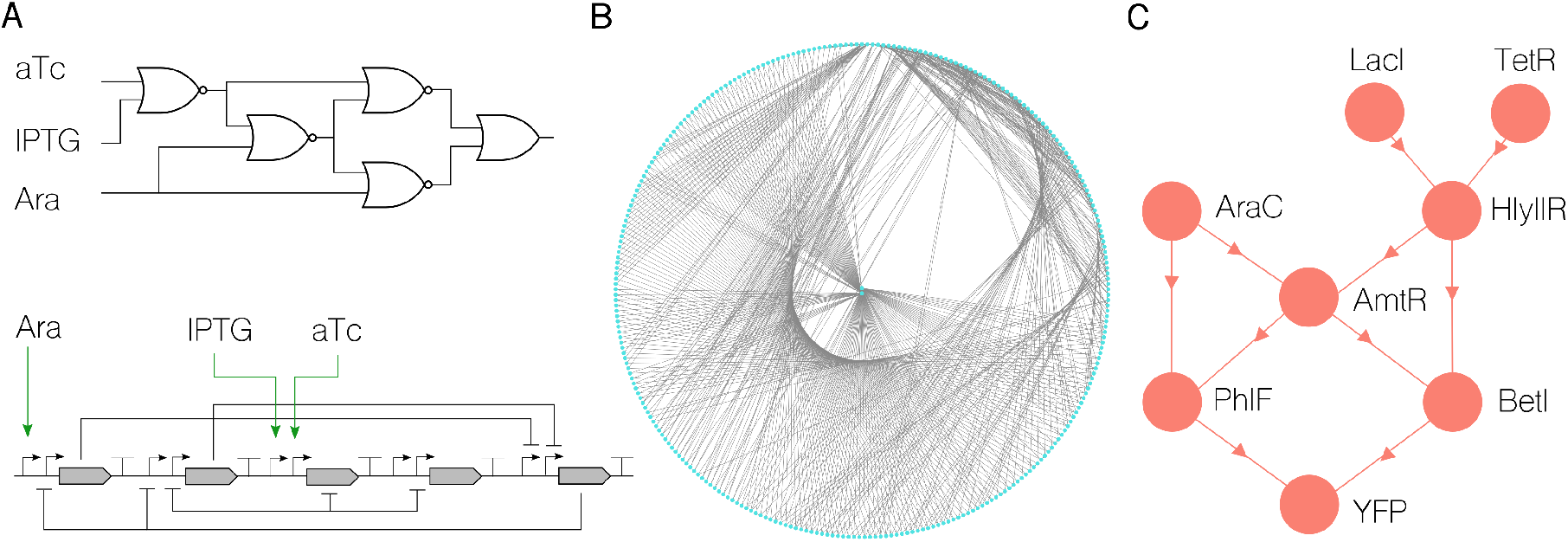
Displaying the protein interaction graph within a complex circuit design. **A** Boolean gene circuit *0x87*^5^ used in this case study. The circuit couples 5 NOR logic gates (top diagram) and uses eight regulatory proteins, five genes and ten promoters (bottom diagram). **B**. Full network with all design information. **C**. Network with only protein (nodes) and interaction (edges) information. Edges represent negative regulation. This view provides a more functional understanding than networks with DNA-only elements.

### Biodesign beyond genetic circuits

The representation of data using networks can be applied to more aspects of synthetic biology away from (but complementary to) circuit designs. In what follows, we briefly cover two such aspects, namely metabolic pathways and experimental protocols, and discuss the potential of networks to provide a general framework for biodesign efforts.

Genetic circuits *run* inside a cellular host (except cell-free systems^45^), which is by no means an austere environment, as the host context, particularly its metabolism, impacts circuit performance, although their interplay is not easy to characterise. A grand challenge is to scale up the complexity of genetic circuits by including metabolic mechanisms that offer dynamics beyond the genetic toolkit catalogue.^18^ As far as the design process is concerned, a question that needs to be answered is whether we can design merged metabolic-genetic circuits. To this end, we show in Figure 5A that the descriptions of a NOR logic gate and a metabolic pathway can dynamically interact if they are encoded into compatible data structures. Specifically, the NOR logic gate uses arabinose as input, which interacts with the same node of the arabinose degradation pathway. Having this information within the same network allows formalising the impact of metabolic dynamics on one of the inputs of the target genetic circuit.

**Figure 5:**
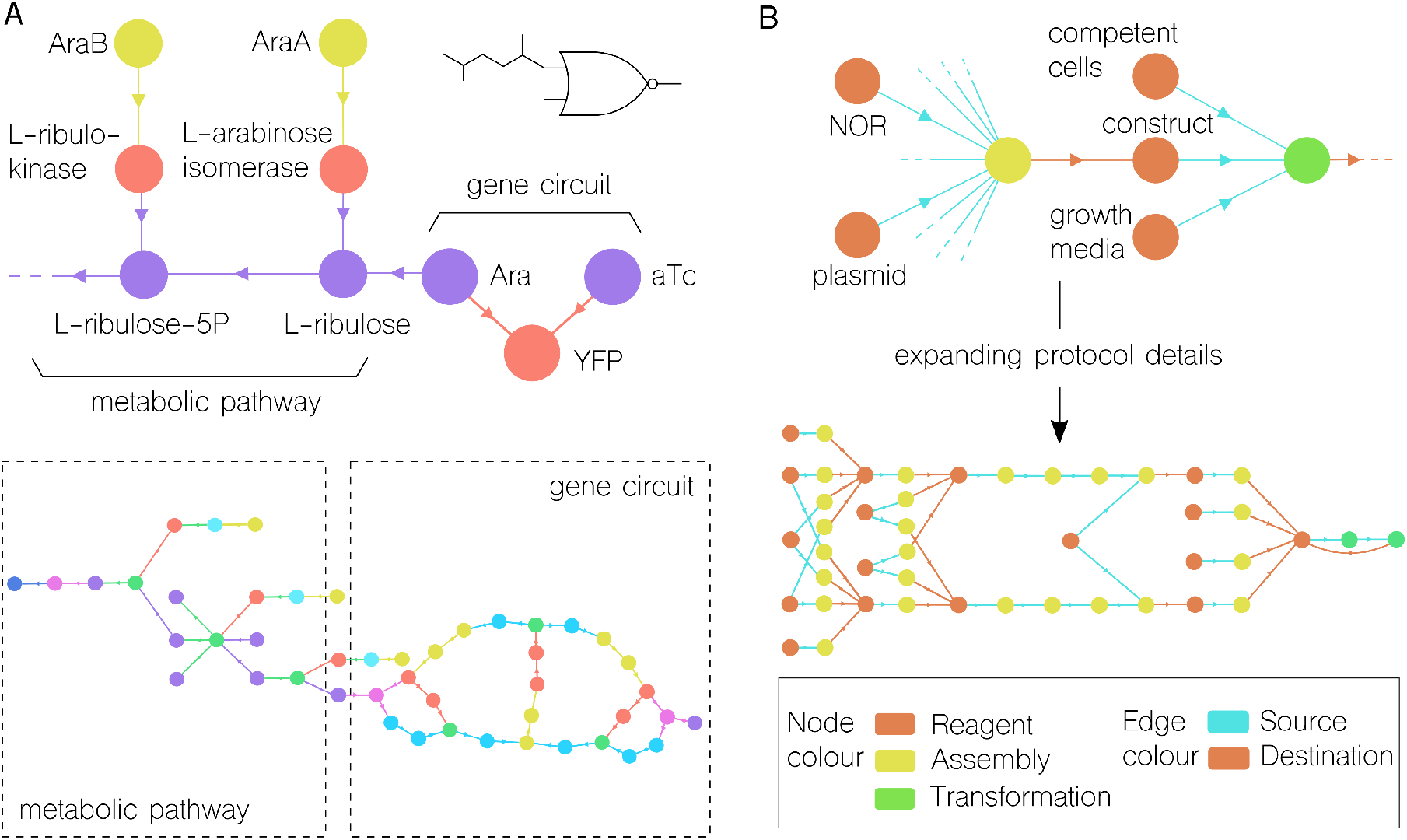
Networks beyond gene circuitry: coupling circuit designs to host metabolic networks and circuit-building protocols. **A**. The network of a gene circuit that uses arabinose as input can interact with the arabinose degradation pathway. Top figure: abstract network displaying critical components of a NOR gate and the initial steps of the arabinose pathway. Bottom figure: linking the corresponding extended networks. **B**. NOR-gate experimental protocol formalized as a network structure. The network can be interactively adjusted to show different levels of abstraction. Nodes represent reagents or sub-protocols, and edges imply input/output relationships.

Finally, we showcase the use of networks for representing experimental protocols. The goal of all circuit designs is to be built and validated experimentally. However, the formalisation of implementation protocols into well-characterised steps and their representation in standard data structures is still a significant challenge^46–48^ that deserves more attention. Figure 5B shows the network that corresponds to the protocol for building and testing the NOR gate used as an example. Here, we chose (from the many options available) to represent materials and methods as nodes and information flow as edges. As in other examples, protocol graphs can also be adjusted at different levels of abstraction. For instance, the assembly node (5B top) includes processes such as restriction, purification and ligation—which are conveniently clustered to provide an overview of the inputs (i.e., what the assembly process gets) and outputs (i.e., what it returns). This network can be linked to the NOR graph at the top node of the hierarchy, therefore having genetic circuits and protocol within the same data structure.

## Conclusion

We have employed a graph-based methodology for representing, visualising, and using circuit design information. Our approach turns design files into networks, which are dynamic structures able to be modified on demand according to user specifications.

When molecular entities, relationships, and other information (e.g. types and roles) are encoded into nodes and edges, a network representation of a genetic design can be established. We have validated this network approach by showcasing its different features, such as abstraction and hierarchy, with structural and functional data within several genetic circuits. The selection of abstraction as a metric to showcase the potential of networks is rooted in the intrinsic complexity of designs and the need to separate high-value information from superfluous details for a given purpose—thus improving understanding for a given individual. We showed that design networks could be automatically adjusted to display different levels of abstraction, from full molecular representation to input/output information only and protein interaction maps. These network manipulations are only an initial subset of possibilities; the vast amount of graph theory methods can directly be applied to design information to analyse networks for many purposes.

The intrinsic modularity of networks allows for coupling genetic circuit designs to other data types providing these are also represented in graphs. We have demonstrated this in two different ways. We showed that a genetic circuit that uses arabinose as input could be automatically coupled to the arabinose degradation pathway graph. By doing this, circuit designs can be extended to include information from their host context, scaling up the functional description of the device. Secondly, we have represented an implementation protocol in network format. While this is just a preliminary effort, which deserves further attention, it shows that protocol networks can also interact with circuit designs for the sake of building a data structure that can be shared along the design-build-test-learn^49^ (DBTL) lifecycle. In short, when data is represented as a graph, merging and clustering potentially disparate entities becomes a far less challenging task, and the graph could be the key to unifying data along the DBTL cycle.

In order to generate high-quality and information-rich networks, designs should capture as much information as possible. Indeed, networks can only work with the provided data—networks cannot fabricate entirely new data, only derived from existing sources. While commonly used formats, such as GenBank, still capture information beyond mere genetic sequences, this information can be challenging to manage computationally due to the inherent informality. Therefore, we advocate using the Synthetic Biology Open Language (SBOL) since it represents formal information, such as modularity or hierarchy, that cannot be captured otherwise. However, our approach does not rely on a specific data format and focuses on representing designs in network structures.

As the complexity of genetic circuits increases, we advocate for networks to manipulate, analyse and communicate design information. We hope networks can maximise the efficiency of design automation procedures and help unification by providing standard^50^ data structures for merged mathematical, genetic, protocol, and other prominent datasets established during synthetic biology projects.

## Methods

### Designs data format and network generation

All design files used in this study were generated using the Synthetic Biology Open Language^29^ (SBOL). While SBOL is defined as Resource Description Framework (RDF) and therefore is graph data, in order to generate the networks discussed here, SBOL designs were converted into an underlying labelled graph model that is more suited for graph theory manipulation. The software package to run that conversion is linked within the *Data and software availability* section below.

### Dynamic network modification

Once a full network is generated from a design, three overlying methods are applied to produce a final user-defined graph. These methods are based on three corresponding preset requirements (namely *view*, *layout* and *attributes*) that respond to the question of what aspects of a design are the focus. The first one (*view*) selects a specific information within a large dataset to provide a predefined perspective. The processes of producing a view include, but are not limited to, pruning unwanted nodes or edges, adding new edges between nodes or abstracting smaller parts of the graph into single nodes to reduce complexity. The second one (*layout*) assists in providing nodes with spatial coordinates, i.e., where nodes are physically positioned relative to one another. The goal of layout adjustments is to minimise the number of edge crossings. The last feature (*attributes*) refer to any visual change to nodes or edges, such as colour, size and shape. Highlighting attributes is essential for comprehension because it allows the addition of extra information without adding new nodes.

### Protocol representation

We used Autoprotocol, a programming language for specifying experimental protocols. Like SBOL, Autoprotocol is captured in a standard format and represented as a graph data structure. We converted protocols into networks, and used the same processes applied to designs, such as defining layouts and other visual additions, for visualising protocol networks.

### Data and software availability

All networks used within Figures have been generated using our own software package that can be accessed from the next repository:https://github.com/intbio-ncl/net_vis_syn_bio. The supplementary data within the repository includes full images of many cutdown networks used within Figures, genetic design files, protocol files and network files containing data to reproduce information views. All data required to replicate the networks described here can be accessed from:https://github.com/MattyCrowther/network-visualisation-supplementary.git. SBOL and Autoprotocol files can be loaded into the software package provided above, while the specific views can be loaded into any tool that accepts common network standards such as Cytoscape and Gephi.

## Acknowledgments

This work was supported by the grants BioSinT-CM (Y2020/TCS-6555) and CONTEXT (Atracción de Talento Program; 2019-T1/BIO-14053) Projects of the Comunidad de Madrid, MULTI-SYSBIO (PID2020-117205GA-I00) and the Severo Ochoa Program for Centres of Excellence in R&D (CEX2020-000999-S) funded by MCIN/AEI /10.13039/501100011033 and the EPSRC studentship 34000024085 (M.C.)

## References

(1) Benenson, Y. Biomolecular computing systems: principles, progress and potential. Nature Reviews Genetics 2012, 13, 455–468.

(2) Brophy, J. A.; Voigt, C. A. Principles of genetic circuit design. Nature methods 2014, 11, 508–520.

(3) Ausländer, S.; Ausländer, D.; Fussenegger, M. Synthetic biology—the synthesis of biology. Angewandte Chemie International Edition 2017, 56, 6396–6419.

(4) Andrianantoandro, E.; Basu, S.; Karig, D. K.; Weiss, R. Synthetic biology: new engineering rules for an emerging discipline. Molecular systems biology 2006, 2, 2006–0028.

(5) Nielsen, A. A.; Der, B. S.; Shin, J.; Vaidyanathan, P.; Paralanov, V.; Strychalski, E. A.; Ross, D.; Densmore, D.; Voigt, C. A. Genetic circuit design automation. Science 2016, 352, aac7341.

(6) Lou, C.; Liu, X.; Ni, M.; Huang, Y.; Huang, Q.; Huang, L.; Jiang, L.; Lu, D.; Wang, M.; Liu, C., et al. Synthesizing a novel genetic sequential logic circuit: a push-on push-off switch. Molecular systems biology 2010, 6, 350.

(7) Friedland, A. E.; Lu, T. K.; Wang, X.; Shi, D.; Church, G.; Collins, J. J. Synthetic gene networks that count. science 2009, 324, 1199–1202.

(8) Shipman, S. L.; Nivala, J.; Macklis, J. D.; Church, G. M. Molecular recordings by directed CRISPR spacer acquisition. Science 2016, 353, aaf1175.

(9) Kawasaki, S.; Ono, H.; Hirosawa, M.; Saito, H. RNA and protein-based nanodevices for mammalian post-transcriptional circuits. Current Opinion in Biotechnology 2020, 63, 99–110.

(10) Wong, A.; Wang, H.; Poh, C. L.; Kitney, R. I. Layering genetic circuits to build a single cell, bacterial half adder. BMC biology 2015, 13, 1–16.

(11) Chen, Y.; Zhang, S.; Young, E. M.; Jones, T. S.; Densmore, D.; Voigt, C. A. Genetic circuit design automation for yeast. Nature Microbiology 2020, 5, 1349–1360.

(12) Lillacci, G.; Benenson, Y.; Khammash, M. Synthetic control systems for high performance gene expression in mammalian cells. Nucleic acids research 2018, 46, 9855–9863.

(13) De Lorenzo, V.; Prather, K. L.; Chen, G.-Q.; O’Day, E.; von Kameke, C.; Oyarzún, D. A.; Hosta-Rigau, L.; Alsafar, H.; Cao, C.; Ji, W., et al. The power of synthetic biology for bioproduction, remediation and pollution control: the UN’s Sustainable Development Goals will inevitably require the application of molecular biology and biotechnology on a global scale. EMBO reports 2018, 19, e45658.

(14) Slomovic, S.; Pardee, K.; Collins, J. J. Synthetic biology devices for in vitro and in vivo diagnostics. Proceedings of the National Academy of Sciences 2015, 112, 14429–14435.

(15) Goñi-Moreno, Á.; Benedetti, I.; Kim, J.; de Lorenzo, V. Deconvolution of gene expression noise into spatial dynamics of transcription factor-promoter interplay. ACS synthetic biology 2017, 6, 1359–1369.

(16) Eldar, A.; Elowitz, M. B. Functional roles for noise in genetic circuits. Nature 2010, 467, 167–173.

(17) Moser, F.; Espah Borujeni, A.; Ghodasara, A. N.; Cameron, E.; Park, Y.; Voigt, C. A. Dynamic control of endogenous metabolism with combinatorial logic circuits. Molecular systems biology 2018, 14, e8605.

(18) Goñi-Moreno, A.; Nikel, P. I. High-performance biocomputing in synthetic biology–integrated transcriptional and metabolic circuits. Frontiers in bioengineering and biotechnology 2019, 7, 40.

(19) Tas, H.; Grozinger, L.; Stoof, R.; de Lorenzo, V.; Goñi-Moreno, A. Contextual dependencies expand the re-usability of genetic inverters. Nature communications 2021, 12, 1–9.

(20) Boo, A.; Ellis, T.; Stan, G.-B. Host-aware synthetic biology. Current Opinion in Systems Biology 2019, 14, 66–72.

(21) Zhu, R.; del Rio-Salgado, J. M.; Garcia-Ojalvo, J.; Elowitz, M. B. Synthetic multistability in mammalian cells. Science 2021, 375, eabg9765.

(22) Grozinger, L.; Amos, M.; Gorochowski, T. E.; Carbonell, P.; Oyarzún, D. A.; Stoof, R.; Fellermann, H.; Zuliani, P.; Tas, H.; Goñi-Moreno, A. Pathways to cellular supremacy in biocomputing. Nature communications 2019, 10, 1–11.

(23) Lopiccolo, A.; Shirt-Ediss, B.; Torelli, E.; Olulana, A. F. A.; Castronovo, M.; Fellermann, H.; Krasnogor, N. A last-in first-out stack data structure implemented in DNA. Nature communications 2021, 12, 1–10.

(24) Appleton, E.; Madsen, C.; Roehner, N.; Densmore, D. Design automation in synthetic biology. Cold Spring Harbor perspectives in biology 2017, 9, a023978.

(25) Kitney, R.; Adeogun, M.; Fujishima, Y.; Goñi-Moreno, Á.; Johnson, R.; Maxon, M.; Steedman, S.; Ward, S.; Winickoff, D.; Philp, J. Enabling the advanced bioeconomy through public policy supporting biofoundries and engineering biology. Trends in biotechnology 2019, 37, 917–920.

(26) Mante, J.; Hao, Y.; Jett, J.; Joshi, U.; Keating, K.; Lu, X.; Nakum, G.; Rodriguez, N. E.; Tang, J.; Terry, L., et al. Synthetic Biology Knowledge System. ACS synthetic biology 2021, 10, 2276–2285.

(27) Misirli, G.; Hallinan, J.; Pocock, M.; Lord, P.; McLaughlin, J. A.; Sauro, H.; Wipat, A. Data integration and mining for synthetic biology design. ACS synthetic biology 2016, 5, 1086–1097.

(28) McLaughlin, J. A.; Myers, C. J.; Zundel, Z.; Misirli, G.; Zhang, M.; Ofiteru, I. D.; Goni-Moreno, A.; Wipat, A. SynBioHub: a standards-enabled design repository for synthetic biology. ACS synthetic biology 2018, 7, 682–688.

(29) Madsen, C.; Moreno, A. G.; Umesh, P.; Palchick, Z.; Roehner, N.; Atallah, C.; Bartley, B.; Choi, K.; Cox, R. S.; Gorochowski, T., et al. Synthetic biology open language (SBOL) version 2.3. Journal of integrative bioinformatics 2019, 16.

(30) Sayers, E. W.; Cavanaugh, M.; Clark, K.; Ostell, J.; Pruitt, K. D.; Karsch-Mizrachi, I. GenBank. Nucleic acids research 2019, 47, D94–D99.

(31) Barabasi, A.-L.; Oltvai, Z. N. Network biology: understanding the cell’s functional organization. Nature reviews genetics 2004, 5, 101–113.

(32) Ideker, T.; Krogan, N. J. Differential network biology. Molecular systems biology 2012, 8, 565.

(33) Krempel, L. Network visualization. The SAGE handbook of social network analysis 2011, 558–577.

(34) Goñi-Moreno, A.; Carcajona, M.; Kim, J.; Martinez-Garcia, E.; Amos, M.; de Lorenzo, V. An implementation-focused bio/algorithmic workflow for synthetic biology. ACS synthetic biology 2016, 5, 1127–1135.

(35) Myers, C. J.; Beal, J.; Gorochowski, T. E.; Kuwahara, H.; Madsen, C.; McLaughlin, J. A.; Misirli, G.; Nguyen, T.; Oberortner, E.; Samineni, M., et al. A standard-enabled workflow for synthetic biology. Biochemical Society Transactions 2017, 45, 793–803.

(36) Beal, J.; Nguyen, T.; Gorochowski, T. E.; Goñi-Moreno, A.; Scott-Brown, J.; McLaughlin, J. A.; Madsen, C.; Aleritsch, B.; Bartley, B.; Bhakta, S., et al. Communicating structure and function in synthetic biology diagrams. ACS synthetic biology 2019, 8, 1818–1825.

(37) Tamsir, A.; Tabor, J. J.; Voigt, C. A. Robust multicellular computing using genetically encoded NOR gates and chemical ‘wires’. Nature 2011, 469, 212–215.

(38) Tas, H.; Grozinger, L.; Goñi-Moreno, A.; de Lorenzo, V. Automated design and implementation of a NOR gate in Pseudomonas putida. Synthetic Biology 2021, 6, ysab024.

(39) Eick, S. Aspects of network visualization. IEEE Computer Graphics and Applications 1996, 16, 69–72.

(40) Karim, R. M.; Kwon, O.-H.; Park, C.; Lee, K. A Study of Colormaps in Network Visualization. Applied Sciences 2019, 9.

(41) Serrano, L. Synthetic biology: promises and challenges. 2007.

(42) Liang, P.; Naik, M. Scaling abstraction refinement via pruning. Proceedings of the 32Nd ACM SIGPLAN Conference on Programming Language Design and Implementation. 2011; pp 590–601.

(43) Calles, B.; Goñi-Moreno, A.; de Lorenzo, V. Digitalizing heterologous gene expression in Gram-negative bacteria with a portable ON/OFF module. Molecular Systems Biology 2019, 15.

(44) Heinemann, M.; Panke, S. Synthetic biology—putting engineering into biology. Bioinformatics 2006, 22, 2790–2799.

(45) Tinafar, A.; Jaenes, K.; Pardee, K. Synthetic biology goes cell-free. BMC biology 2019, 17, 1–14.

(46) Yaman, F.; Adler, A.; Beal, J. AI challenges in synthetic biology engineering. Proceedings of the AAAI conference on artificial intelligence. 2018.

(47) Beal, J.; Weiss, R.; Densmore, D.; Adler, A.; Appleton, E.; Babb, J.; Bhatia, S.; Davidsohn, N.; Haddock, T.; Loyall, J., et al. An end-to-end workflow for engineering of biological networks from high-level specifications. ACS Synthetic Biology 2012, 1, 317–331.

(48) Sainz de Murieta, I.; Bultelle, M.; Kitney, R. I. Toward the first data acquisition standard in synthetic biology. ACS synthetic biology 2016, 5, 817–826.

(49) Tellechea-Luzardo, J.; Otero-Muras, I.; Goñi-Moreno, A.; Carbonell, P. Fast biofoundries: coping with the challenges of biomanufacturing. Trends in Biotechnology 2022,

(50) Beal, J.; Goñi-Moreno, A.; Myers, C.; Hecht, A.; de Vicente, M. d. C.; Parco, M.; Schmidt, M.; Timmis, K.; Baldwin, G.; Friedrichs, S., et al. The long journey towards standards for engineering biosystems: Are the Molecular Biology and the Biotech communities ready to standardise? EMBO reports 2020, 21, e50521.

